# A stickiness scale for disordered proteins

**DOI:** 10.64898/2026.01.25.701651

**Authors:** Fan Cao, Giulio Tesei, Kresten Lindorff-Larsen

## Abstract

Disordered proteins are a heterogeneous group of proteins that play a broad range of functions in biology, and display conformational properties that range from compact globules to expanded chains. We here describe the results of a data-driven approach to derive a scale that represents the propensity of the twenty amino acids to interact with one another relative to water. The scale is based on biophysical experiments on 115 proteins and can be thought of as a ‘stickiness’ (or hydropathy) scale specific for disordered proteins. We compare the scale to 70 other previously reported hydropathy scales and find that it is closer to four scales related to membrane proteins or the transition temperatures of elastin-like peptides. We envisage that the new scale will be useful in bioinformatics and machine learning approaches to quantify the role of sequence composition and patterning in disordered proteins, to understand the driving forces for their interactions with other molecules, and their evolutionary conservation.

## Introduction

Disordered proteins are a heterogeneous group of proteins that carry out a broad range of functions in biology (***Holehouse and Kragelund, 2024***). As a group, one shared characteristic is that they display a substantial amount of conformational disorder without fixed secondary and tertiary structure; this conformational flexibility in turn underlies the myriad functions to which disordered proteins contribute in a cellular context.

While the functions of disordered proteins are many and varied, several principles have emerged as to how their functions are encoded in their sequences. One common theme is that, in many cases, the overall polymeric properties of the ensemble of conformations adopted by a protein chain, such as its degree of expansion, can play a key role in the interactions and functions of the disordered protein (***Holehouse and Alberti, 2025***). Combinations of theory, simulations, and experiments have shed light on how the amino acid sequence of a disordered protein determines its conformational properties, and have highlighted how the overall composition and patterning of amino acids, rather than the exact location of each amino acid, is key to determine ensemble properties (***Das et al., 2015; Tesei et al., 2024; Lotthammer et al., 2024***).

One key parameter of sequence–ensemble relationships for disordered proteins is how hydrophobic residues are present and distributed throughout the polypeptide chain. Hydrophobicity is, however, a complex phenomenon and there are many different ways to probe and quantify it. In the context of disordered proteins it has been shown that aromatic residues can have a particular strong effect on conformational properties (***Martin et al., 2020***), possibly also due to their ability to form *π*–*π* interactions (***Vernon et al., 2018***). There are, however, many different hydrophobicity scales that have been measured over the years (***Simm et al., 2016; Ji et al., 2023; Lobo et al., 2025***), and it is not fully clear which of these—if any—are appropriate for an accurate description of the conformational properties of disordered proteins (***Tesei et al., 2021b***).

In recent years, there has been considerable interest in defining appropriate hydropathy scales for the 20 common amino acids in the context of disordered proteins. Such scales can, for example, be used as input to construct coarse-grained simulation models for disordered proteins (***Dignon et al., 2018; Regy et al., 2021***), or to relate the hydrophobicity patterning of the linear amino acid sequence to conformational properties (***Zheng et al., 2020***). However, these hydropathy scales vary considerably and give substantially different descriptions of the conformational properties of disordered proteins (***Dignon et al., 2018; Regy et al., 2021***).

We and others have taken a different approach to parameterizing coarse-grained simulation models by learning the appropriate scale directly from experimental data that describe the conformational properties of disordered proteins (***Norgaard et al., 2008; Demerdash et al., 2019; Latham and Zhang, 2019; Martin et al., 2020; Tesei et al., 2021b; Mugnai et al., 2025***). Specifically, we have used a Bayesian parameter optimization scheme (***Norgaard et al., 2008***) to learn a set of 20 parameters in a molecular force field that can be used to predict conformational ensembles of disordered proteins. We have applied this approach to learn a relative ‘stickiness’ scale for amino acids in our CALVADOS (Coarse-graining Approach to Liquid-liquid phase separation Via an Automated Datadriven Optimisation Scheme) force field (***Tesei et al., 2021b; Tesei and Lindorff-Larsen, 2023***). This force field parameterizes the energetics in a one-bead-per-residue model via contributions from bonded terms and non-bonded terms that model ionic (electrostatic) interactions and other interactions such as those driven by hydrophobicity. The latter term is parameterized via 20 so-called *λ* values that are combined to estimate pair-wise van der Waals interactions between amino acid types. While there are 210 possible different pairs of amino acid types, studies of folded and disordered proteins have suggested that it is possible to represent these different interactions relatively accurately in terms of one parameter for each amino acid that can then be combined to represent the full set of pairs (***Cieplak et al., 2001; Dannenhoffer-Lafage and Best, 2021***).

In the context of the CALVADOS model we have shown that it is possible to fit experimental nuclear magnetic resonance (NMR) and small-angle X-ray scattering (SAXS) data reporting on the global structural properties of disordered proteins by fitting 20 *λ* values (one for each amino acid type) (***Tesei et al., 2021b; Tesei and Lindorff-Larsen, 2023***). The resulting parameters resonate with previous analyses scoring the importance of different types of interactions in disordered proteins (***Martin et al., 2020; Bremer et al., 2022***). We also compared the CALVADOS *λ* values to a range of different hydropathy scales (***Simm et al., 2016***) and found them to be in relatively good agreement with, for example, the Urry scale (***Urry et al., 1992; Tesei et al., 2021b***), which was independently selected as an alternative scale for use in simulations of disordered proteins (***Regy et al., 2021***). These results suggest that our parameter learning scheme is useful to define a scale for the physicochemical interactions within and between disordered proteins.

In both the CALVADOS and related models, the interactions are, however, defined by other parameters than the 20 *λ* values. Of particular relevance to interpreting the physical origins of the *λ* values is that the Ashbaugh-Hatch potential describing non-ionic, non-bonded interactions is parameterized by the combination of the *λ* value with the diameter of the amino acid bead, *σ*, that determines the range of interaction (***Ashbaugh and Hatch, 2008***). Thus, while the *λ* values of, for example, glycine (*λ* = 0.706) is similar to that of arginine (*λ* = 0.731), arginine is a more ‘sticky’ residue in the context of the CALVADOS model because its range of interaction is greater than that of glycine (*σ* = 0.656 nm vs. *σ* = 0.45 nm). Similarly, tyrosine and tryptophan have similar *λ* values in the CALVADOS 2 model, but tryptophan forms stronger interactions because its interactions are longer-ranged (*σ* = 0.678 nm vs. *σ* = 0.646 nm). The variation in *σ* thus makes it difficult to compare the relative stickiness of the 20 amino acids from the *λ* values because they also have different *σ* values.

Here, we describe a new stickiness scale for the 20 common amino acids in the context of disordered proteins. We optimize these in the context of the CALVADOS model using experimental data reporting on the compaction and long-range interactions within disordered proteins. In contrast to previous work we optimize these using a single, common interaction range for all amino acids. In this way, the resulting *λ* values contain the joint contributions from both variation in the size and relative strength of interactions of the different amino acids. We find that the resulting scale conforms to the expected pattern based on previous analyses and highlights the unique character of the interactions within and between disordered regions in proteins.

## Results and Discussion

We used our previously described parameter-learning procedure (***Norgaard et al., 2008; Tesei et al., 2021b; Tesei and Lindorff-Larsen, 2023***) to derive stickiness scales that result in accurate predictions of conformational properties in the context of a coarse-grained simulation model of disordered regions and proteins. As a training set, we collected radii of gyration (*R*_g_) from SAXS experiments for 115 disordered proteins. These include 76 proteins that we used for previous work (***Tesei and Lindorff-Larsen, 2023; Cao et al., 2024***), 23 additional proteins that we previously used for validation (***Tesei et al., 2024***), and data for 16 proteins that we collected from the literature for this study (Tab. S1). We computed the frequency of occurrence of each amino acid in both the new training set and the previously used dataset. We observed that the fraction of most amino acids has changed only slightly (Fig. 1).

**Figure 1.**
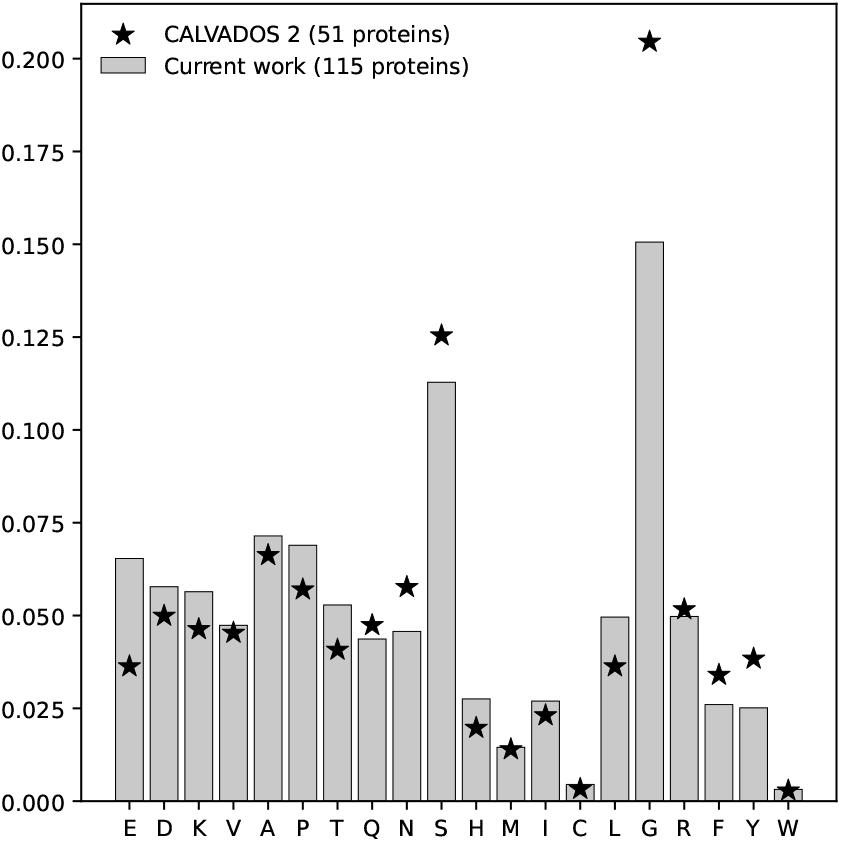
Fraction of amino acids in the training set of CALVADOS 2 (stars) and in the expanded training set used in this work (bars).

We made minor modifications to our optimization algorithm (Methods) including using either a single, shared interaction range (⟨*σ*⟩ = 0.56 nm) for all 20 residue types, or the residue-specific *σ*_AA_ values (***Kim and Hummer, 2008***) that we used to parameterize CALVADOS (***Tesei et al., 2021b***). While using the shared ⟨*σ*⟩ results in a less physically realistic simulation model, this choice ensures that the resulting 20 *λ* values reflect the effective stickiness of the 20 amino acids without the complications of interpreting the combined effect of changes in both *λ* and *σ* on the strength and distance dependence of the interactions. In addition to the training set, we compiled a validation dataset consisting of 20 proteins with SAXS data, 4 proteins with paramagnetic relaxation enhancement (PRE) NMR data, and 22 proteins with Förster resonance energy transfer (FRET) data to validate the resulting models (Tabs. S2, S3 and S4).

With this setup, we performed the optimization and validation procedures in the following steps:

1. We used our previously described force field parameterization approach (***Norgaard et al., 2008; Tesei et al., 2021b***) to learn *λ* values that minimize the deviation between experimental and simulated *R*_g_ values (Eq. 1). To avoid overfitting on the training set, we introduced a regularization term, *θ*MSE (*λ*_CALVADOS2_, *λ*_trial_ ), that penalizes deviations between CALVADOS 2 and new *λ* values (Methods). The strength of the regularization term is determined by the hyperparameter *θ*.
2. To examine the robustness regarding the choice of *θ*, we performed optimization runs using *θ* values ranging between 150 and 4,800, using the residue-specific *σ*_AA_ values. For each resulting model, we quantified the agreement with the validation data set as 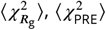, and 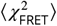 (Eqs. 2, 3, and 4).
3. Since we use simulated annealing with Monte Carlo sampling, repeated optimization runs yield slightly different parameters. Therefore, for each *θ* value, we performed three independent optimizations and reported means and standard error of the mean (SEM) for 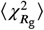 evaluated on the training set (Fig. 2A) as well as 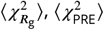, and 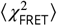 evaluated on the vali-dation sets (Fig. 2B-D). 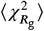 evaluated on the training set increases with increasing *θ* (Fig. 2A) whereas the agreement with the validation sets shows smaller variations (Fig. 2B-D). When *θ* is small, the optimization is mainly driven by 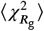 leading to overfitting and limiting the transferability of the resulting model, as reflected by the larger ⟨*χ*^2^⟩ values across the validation sets (Fig. 2B-D). We note that since we use the CALVADOS 2 *λ* values as prior, the results are relatively good even at high values of *θ*. For *θ* ≈ 1,950, we reach good balance between 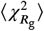 and MSE *λ*_CALVADOS2_, *λ*_trial_, as indicated by the local minima in ⟨*χ*^2^⟩ across the validation sets. Further increasing *θ* results in underfitting and higher ⟨*χ*^2^⟩ values on the validation sets.
4. With *θ* = 1, 950, we performed three additional independent optimization runs, this time using the average ⟨*σ*⟩, and calculated means and SEM for the ⟨*χ*^2^⟩ alues across the same training and validation sets as for the optimizations with residue-specific *σ*_AA_ values.

**Figure 2.**
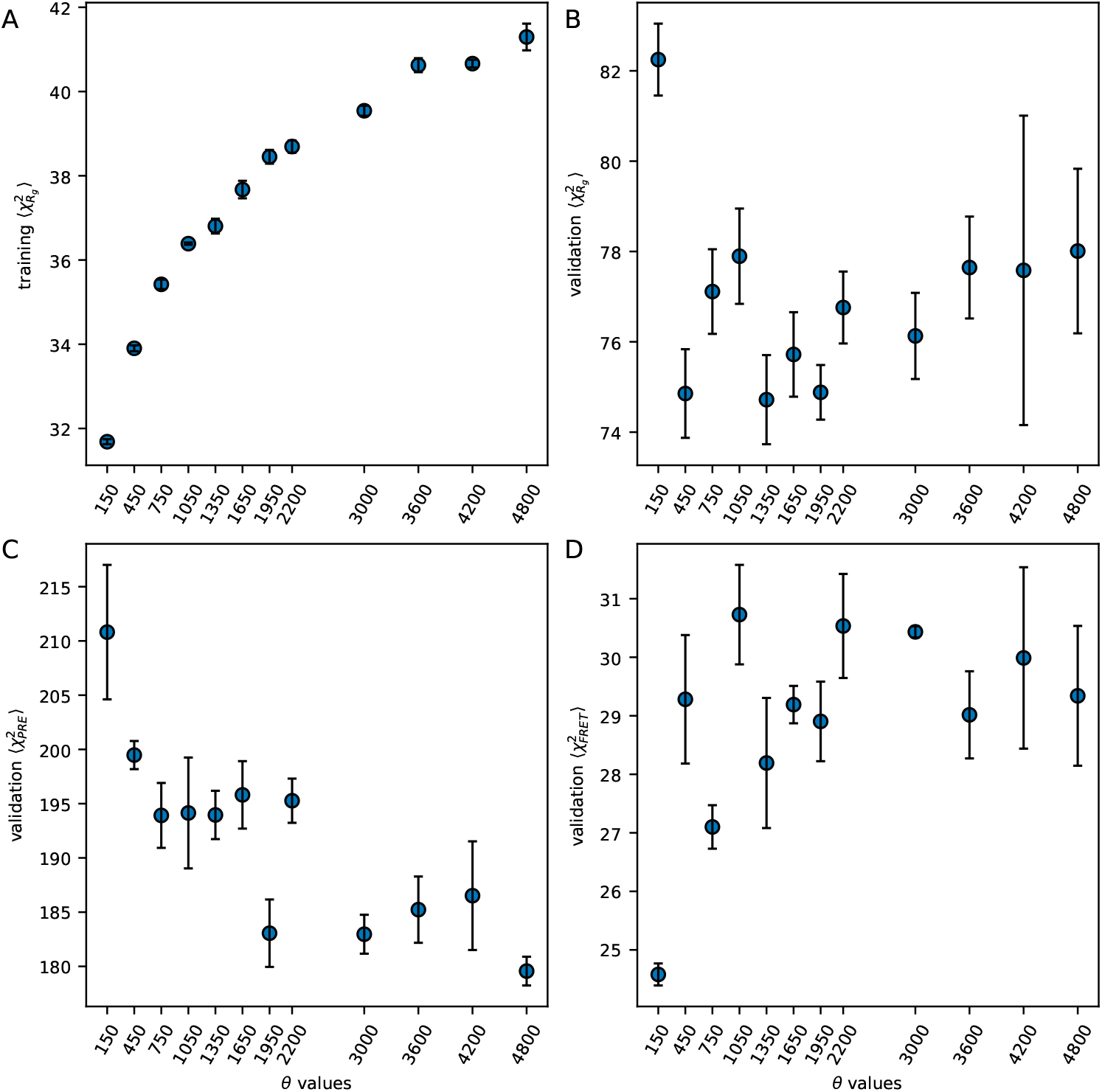
Optimization and cross-validation results for the model with residue-specific *σ*_AA_ values.(A) 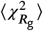 evaluated on the training set as a function of the *θ* value weighting the regularization term. (B–D) ⟨*χ*^2^⟩ values evaluated on the SAXS (B), PRE NMR (C), and FRET (D) validation sets with varying *θ* values. Error bars are SEM reported as 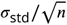 where *σ*_std_ is the standard deviation over three independent optimization runs and *n* is the number of independent optimization runs.

The two resulting sets of *λ* values (Fig. 3A and Tab. S5) are overall similar to each other (Pearson correlation coefficient of 0.99) and to both CALVADOS 2 (***Tesei and Lindorff-Larsen, 2023***) and CALVADOS 3 (***Cao et al., 2024***) *λ* values. There are, however, some notable differences that result from optimizing either in the context of a single or variable diameter. For the smallest and most abundant amino acid in the training set, glycine, there is a drop in *λ* in the ⟨*σ*⟩ model (Fig. 3B). This suggests that the lower *λ* compensates for the relatively longer interaction range when glycine is represented with a larger diameter. We also observed a similar drop in *λ* for serine, which is the second most abundant amino acid in our training set (Fig. 3B). Similarly, the *λ* values of valine, methionine, and leucine increase when these residue types are represented with the smaller average ⟨*σ*⟩ values. However, this compensation of an increase in *λ* by a decrease in *σ*, and vice versa, is not observed consistently for the remaining amino acid types (Fig. 3B).

**Figure 3.**
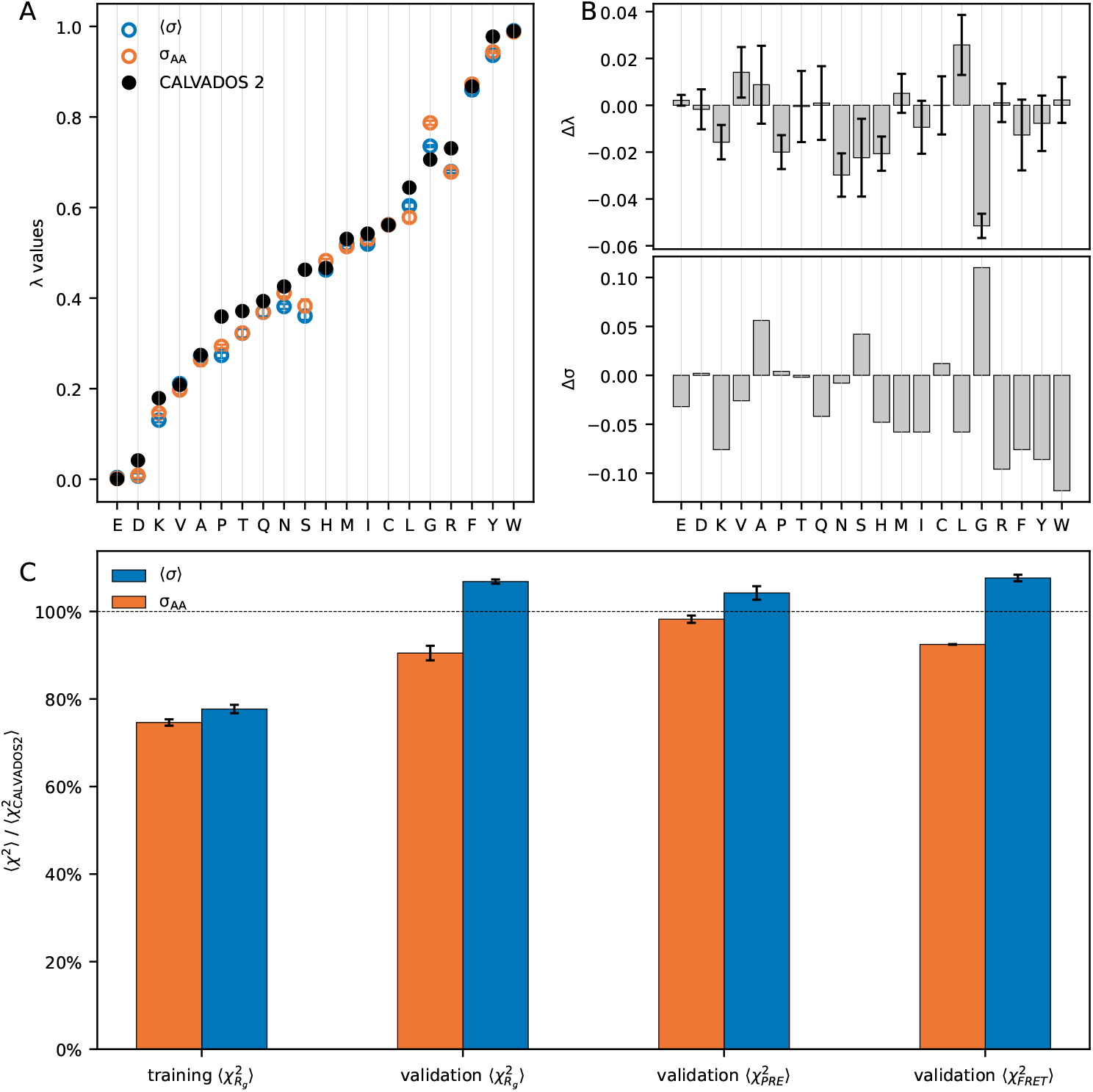
Optimized stickiness scales for disordered proteins using *θ* = 1, 950. (A) We show the 20 *λ* values that we optimized using either a single diameter shared for all 20 amino acids (⟨*σ*⟩ = 0.56 nm, blue), or residue-specific diameters (*σ*_AA_, orange). CALVADOS 2 *λ* values are shown for comparison (black). Error bars are standard deviations over three independent optimization runs. (B) Differences in *λ* and *σ* of two optimization setups (*λ* and *σ* values in ⟨*σ*⟩ minus *λ* and *σ* values in *σ*_AA_, respectively). We use the difference between the averaged lambdas of two setups over three independent optimizations to compare. Error bars are 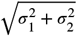, where *σ*_1_ and *σ*_2_ are the standard deviations of two setups over three independent optimization runs. (C) Comparison of ⟨*χ*^2^⟩ obtained with average (⟨*σ*⟩ = 0.56 nm, blue) and residue-specific diameters (*σ*_AA_, orange) normalized by the corresponding values for the original CALVADOS 2 model, 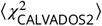 Error bars are SEM reported as 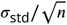, where *σ*_std_ is the standard deviation over three independent optimization runs and *n* is the number of independent optimization runs.

We found that the optimizations performed with average and variable diameters resulted in models that agree similarly well with the experimental SAXS data in the training set (Fig. 3C). We noted that relative mean signed deviations, rMSD = ⟨ (*R*_g,sim_ − *R*_g,exp_)/*R*_g,exp_⟩ (which probes any bias towards over- or under-estimating radii of gyration) for the residue-specific *σ*_AA_ and shared ⟨*σ*⟩ (−0.008 and −0.012, respectively) are considerably lower than the rMSD for CALVADOS 2 (−0.029), which was optimized against only a subset of the current training set (Fig. 4).

**Figure 4.**
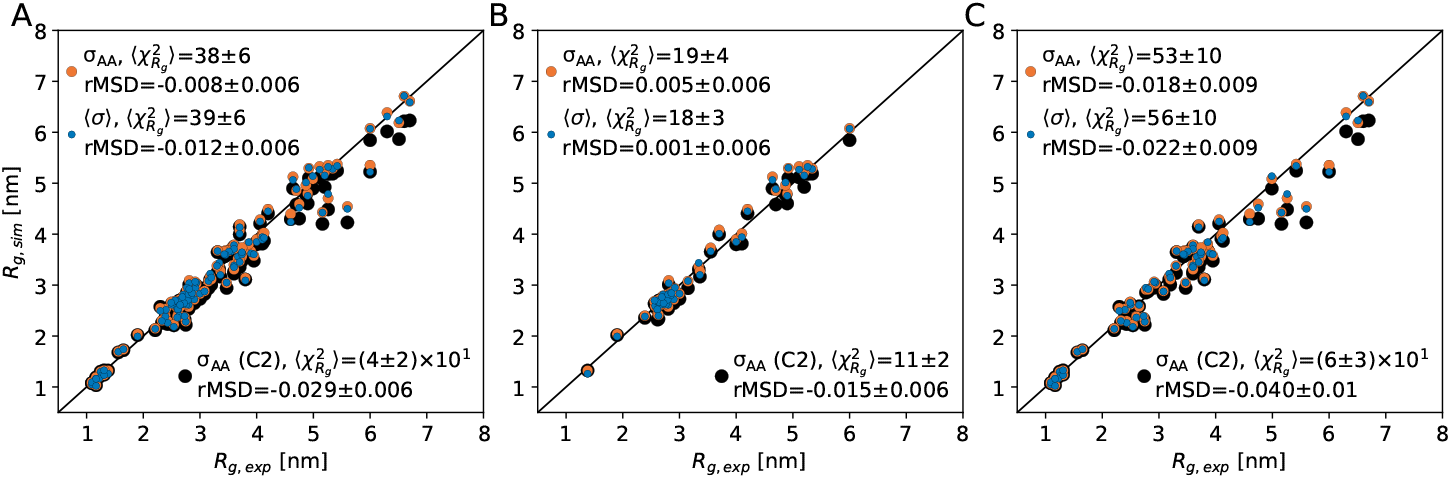
Performance of the models derived from the residue-specific *σ*_AA_ (orange), the shared ⟨*σ*⟩ (blue) and CALVADOS 2 (black) on (A) the whole training set, (B) the training set for CALVADOS 2 and (C) the newly added training data. Relative mean signed deviations, rMSD = ⟨ (*R*_g,sim_ − *R*_g,exp_)/*R*_g,exp_⟩, are reported in the legends with the SEM calculated using bootstrapping. A negative rMSD value indicates that the calculated radii of gyration are systematically lower than the experimental values.

The validation of the two sets of parameters obtained with *σ*_AA_ and ⟨*σ*⟩ instead shows that the model derived using a single average diameter is less transferable and performs considerably worse than CALVADOS 2 whereas the model with variable diameters outperforms CALVADOS 2 across all training and validation sets (Fig. 3C). This suggests that one should use the combination of *σ*_AA_ and its resulting *λ* values to simulate disordered proteins for more accurate conformational ensembles, rather than using ⟨*σ*⟩ and the corresponding *λ* values. The latter values can instead be used as a scale for example in bioinformatics analyses.

With new stickiness scales for disordered proteins in hand, we then asked which of the many other scales they are most similar to. We therefore used hierarchical clustering to construct a dendrogram of the new scales, together with those from the CALVADOS 2 and HPS-Urry models (***Urry et al., 1992; Regy et al., 2021***), and 70 hydrophobicity scales that we collected from the literature (***Simm et al., 2016***) and previously used as prior to develop CALVADOS 2 (***Tesei and Lindorff-Larsen, 2023***). The results show that the new scales are most similar to the CALVADOS 2 scale and a cluster comprising the Ponnuswamy-and-Gromiha scale from crystal structures of membrane proteins (***Ponnuswamy and GROMIHA, 1993***), the Urry scale from transition temperatures of elastin-like peptides (***Urry et al., 1992***), the Wimley-and-White scale from peptides partitioning at the lipid bilayer-water interface (***Wimley and White, 1996***) and the related Bishop scale (***Bishop et al., 2001***) (Fig. 5). The agreement with the CALVADOS 2 scale is not surprising given that this was used as a prior in the optimization, and we noted that a scale related to the Urry scale has previously also been shown to work well for simulations of disordered proteins (***Regy et al., 2021***).

**Figure 5.**
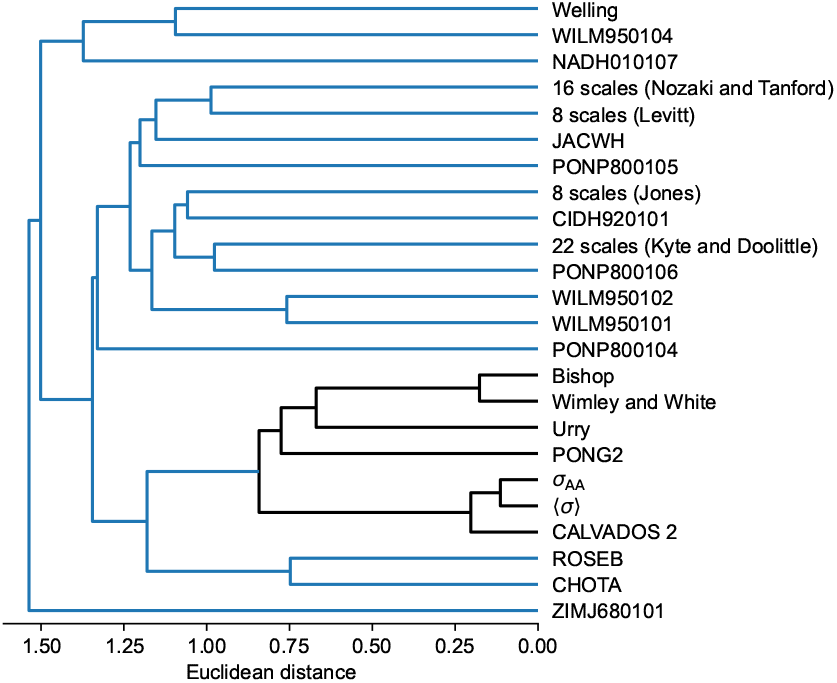
Dendrogram of a hierarchical clustering of 70 hydrophobicity scales from the literature, the CALVADOS 2 stickiness scale and the new scales presented in this work (*σ*_AA_ and ⟨*σ*⟩ ). The clustering was based on Euclidean distances between the scales after min-max normalization.

## Conclusions

Disordered proteins play a wide range of biological functions that may be related to their global structural and polymer properties. These properties are determined by an interplay between sequence composition and patterning, and involve interactions between hydrophobic and charged residues and solubilizing effects of more polar amino acids—all in balance with solvent interactions. Here, we have presented a new hydropathy (‘stickiness’) scale for disordered proteins with the aim of gaining a better understanding of sequence–ensemble–function relationships. The scale may for example be used as features to predict biophysical properties from sequence (***Zheng et al., 2020; Tesei et al., 2024; von Bülow et al., 2025***) or used to help represent sequences in alignment free approaches (***Zarin et al., 2019; Singleton and Eisen, 2024; Chow et al., 2024; Ruff et al., 2025; Halpin and Keating, 2025***). We derive this scale using a Bayesian parameter-learning scheme from biophysical data on 115 disordered proteins. In contrast to previous scales that conflate stickiness and size of the amino acids, we have derived this scale in such a way that interaction strength and distance-dependence scale are encoded in a single number. We hope that the scale will be useful, for example, for predicting conformational and biophysical properties of disordered proteins including their ability to self-associate and phase separate.

## Methods

### Experimental data

We used the previously described CALVADOS model with the *λ* and *σ* parameters specified in this work to perform coarse-grained simulations (***Tesei et al., 2021b; Tesei and Lindorff-Larsen, 2023***). We generated initial conformations of all disordered proteins as Archimedes’ spirals with a distance of 0.38 nm between connecting beads. We then conducted Langevin dynamics simulations using OpenMM 7.6.0 (***Eastman et al., 2017***) in the NVT ensemble with an integration time step of 10 fs and friction coefficient of 0.01 ps^−1^. Single chains of disordered proteins with *N* residues were simulated in a cubic box with a (*N* − 1)×0.38 + 4 nm box edge length under periodic boundary conditions. Each disordered protein was simulated in 20 replicas for 6.3 *ns*∼139 *ns* depending on the sequence length (***Tesei and Lindorff-Larsen, 2023; Tesei et al., 2024***). Final trajectories had 4,000 frames for each protein, discarding the initial 10 frames in each replica for equilibration.

### Parameter optimization

To optimize the stickiness parameters, *λ*, we used a Bayesian parameter-learning procedure (***Tesei and Lindorff-Larsen, 2023***) which we modified here to minimize the following loss function:

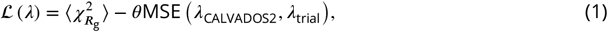

where 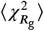 quantifies the *R*_g_ deviations between experiments and simulations over all sequences in the training set,

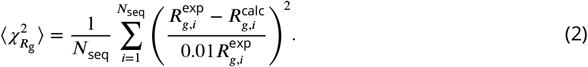

Here, MSE 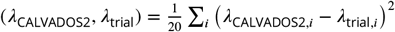 is the mean square error between *λ* values of CALVADOS 2 and trial *λ* values. In a Bayesian framework, the MSE term corresponds to assuming normal distributions centered around *λ*_CALVADOS2,*i*_ as prior. We used 1% of the experimental *R*_g_ values as the errors to calculate 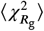 in Eq. 2. We started the optimization using CALVADOS 2 as the initial *λ* values and we found that the optimization converged within the first cycle. Therefore, we selected the *λ* values with the lowest loss during the first cycle of optimization to validate resulting models on the validation set. During optimization, we restrict *λ* to positive values while we allow it to be greater than one.

We screened *θ* from 150 to 4,800 in order to select a *θ* value to balance 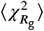 and MSE(*λ*_CALVADOS2_, *λ*_trial_). We performed three independent optimization runs for each *θ* value. We then evaluated the agreement with the validation set for each resulting parameter set (Fig. 2, Tab. S2, Tab. S3 and Tab. S4). Based on cross validation results, we chose *θ* = 1, 950 as the prior weight to avoid overfitting. We used Eq. 2 to calculate 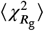 of the proteins with SAXS data in the validation set, and used the following equation to calculate 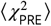:

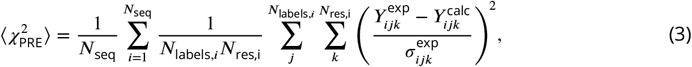

where *N*_labels,*i*_ and *N*_res,*i*_ are the numbers of spin-labeled mutants and measured residues on sequence *i*, respectively, 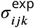 is the experimental error, *Y* is either *I*_para_/*I*_dia_ or Γ_2_ calculated using the rotamer library approach implemented in DEER-PREdict (***Tesei et al., 2021a***). We used the equation below to calculate 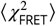:

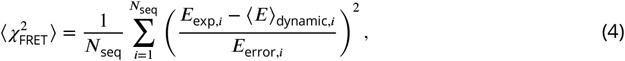

where *E*_exp,*i*_ is the experimental FRET efficiency and *E*_error,*i*_ is the experimental error. ⟨*E*⟩ _dynamic,*i*_ is the calculated FRET efficiency and written as (***Montepietra et al., 2024***):

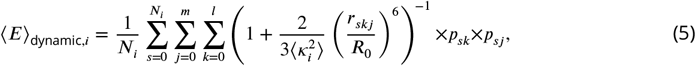

Where 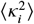 is calculated over all the protein conformations of sequence *i* and combinations of probe rotamers as

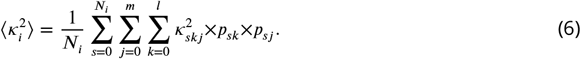

Here, *N*_*i*_, and are the numbers of protein conformations of sequence *i*, rotamers for the donor and the acceptor, respectively, *p*_*sk*_ and *p*_*sj*_ are weights corresponding to the rotamers *k* and *j* of the fluorophores in conformation 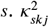 is an instantaneous value calculated for a given combination of donor and acceptor rotamers. *R*_0_ is Förster radius, and *r*_*skj*_ is the donor–acceptor distance (***Montepietra et al., 2024***).

### Clustering method

We used the 70 hydrophobicity scales from Tables 3 and 4 of Simm et al. (***Simm et al., 2016***), previously selected to be non-identical after min-max normalization (***Tesei and Lindorff-Larsen, 2023***). Together with the CALVADOS 2 stickiness scale and the new scales introduced in this work, the normalized scales were clustered using hierarchical agglomerative clustering based on Euclidean distances and the average linkage method, as implemented in scikit-learn v1.0.2 (***Pedregosa et al., 2011***).

## Data and software availability

Scripts and data to reproduce the work are available via https://github.com/KULL-Centre/_2026_cao_scales.

## Acknowledgments

We acknowledge the use of computational resources from Computerome 2.0, the ROBUST Resource for Biomolecular Simulations (supported by the Novo Nordisk Foundation; NNF18OC0032608) and the Biocomputing Core Facility at the Department of Biology, University of Copenhagen. This research was supported by the PRISM (Protein Interactions and Stability in Medicine and Genomics) centre funded by the Novo Nordisk Foundation (NNF18OC0033950, to K.L.-L.), and CSC (China scholarship council, 202206340019).

---

**Table S1.** Experimental solution conditions and radii of gyration of the disordered proteins included in the training set for the Bayesian parameter-learning procedure. Errors of experimental *R*_g_ are estimated as 1% of *R*_g_ values.

| Protein | N | $R_g$ [nm] | T [K] | $c_s$ [M] | pH | Ref. |
| --- | --- | --- | --- | --- | --- | --- |
| S130_143 | 16 | 1.093 | 298.15 | 0.161 | 8.0 | (Koren et al., 2023) |
| S66_81 | 17 | 1.168 | 298.15 | 0.161 | 8.0 | (Koren et al., 2023) |
| S110_125 | 18 | 1.164 | 298.15 | 0.161 | 8.0 | (Koren et al., 2023) |
| S82_96 | 18 | 1.168 | 298.15 | 0.161 | 8.0 | (Koren et al., 2023) |
| S45_64 | 20 | 1.221 | 298.15 | 0.161 | 8.0 | (Koren et al., 2023) |
| S106_128 | 20 | 1.248 | 298.15 | 0.161 | 8.0 | (Koren et al., 2023) |
| S87_105 | 20 | 1.309 | 298.15 | 0.161 | 8.0 | (Koren et al., 2023) |
| S67_86 | 20 | 1.31 | 298.15 | 0.161 | 8.0 | (Koren et al., 2023) |
| S26_45 | 20 | 1.094 | 298.15 | 0.161 | 8.0 | (Koren et al., 2023) |
| S129_146 | 21 | 1.267 | 298.15 | 0.161 | 8.0 | (Koren et al., 2023) |
| Hst5 | 24 | 1.38 | 298.0 | 0.168 | 7.0 | (Jephthah et al., 2019) |
| FCP1 | 32 | 1.56 | 298.0 | 0.11 | 6.5 | (Gibbs and Showalter, 2016) |
| Hst52 | 48 | 1.9 | 298.0 | 0.168 | 7.0 | (Fagerberg et al., 2020) |
| TRF2_BR | 49 | 1.65 | 277.0 | 0.15 | 7.0 | (Nečasová et al., 2017) |
| MenVL | 62 | 2.54 | 283.0 | 0.15 | 8.5 | (Webby et al., 2021) |
| p532070 | 62 | 2.39 | 277.0 | 0.1 | 7.0 | (Zhao et al., 2021) |
| DomainV | 67 | 2.43 | 288.15 | 0.1985 | 7.0 | (Chan-Yao-Chong et al., 2019) |
| ACTR | 71 | 2.63 | 278.0 | 0.2 | 7.4 | (Kjaergaard et al., 2010) |
| DSS1 | 71 | 2.5 | 288.0 | 0.17 | 7.4 | (Pesce et al., 2023) |
| GRB14_PIR | 74 | 2.6 | 293.0 | 0.13 | 8.0 | (Moncoq et al., 2004) |
| HeV_PNT3_CTD | 77 | 2.75 | 293.0 | 0.12 | 7.2 | (Nilsson et al., 2022) |
| HeV_PNT3_CTD_3Y3A | 77 | 2.73 | 293.0 | 0.12 | 7.2 | (Nilsson et al., 2022) |
| Ash1 | 81 | 2.9 | 293.0 | 0.15 | 7.5 | (Martin et al., 2016; Jin and Gräter, 2016) |
| CTD2 | 83 | 2.614 | 293.0 | 0.12 | 7.5 | (Jin and Gräter, 2021; Gibbs et al., 2019) |
| Sic1 | 92 | 3.0 | 293.0 | 0.2 | 7.5 | (Gomes et al., 2020) |
| sfAFP | 92 | 2.3 | 293.15 | 0.351 | 7.5 | (Baxa et al., 2024) |
| SH4UD | 95 | 2.71 | 293.15 | 0.216 | 8.0 | (Shrestha et al., 2019) |
| BMAL1P624A | 98 | 2.77 | 283.25 | 0.154 | 7.2 | (Garg et al., 2019) |
| CoINT | 98 | 2.8 | 277.0 | 0.433 | 7.6 | (Johnson et al., 2017) |
| VWF | 103 | 3.08 | 293.0 | 0.153 | 7.4 | (del Amo-Maestro et al., 2021) |
| Fez1 | 103 | 3.6 | 293.0 | 0.168 | 7.4 | (Alborghetti et al., 2010) |
| UL11 | 104 | 2.4 | 277.0 | 0.11 | 7.5 | (Metrick et al., 2020) |
| lambdaN | 107 | 3.8 | 293.0 | 0.11 | 7.0 | (Johansen et al., 2011) |
| p27Cv78 | 107 | 2.211 | 293.0 | 0.095 | 7.2 | (Das et al., 2016) |
| p27Cv56 | 107 | 2.328 | 293.0 | 0.095 | 7.2 | (Das et al., 2016) |
| p27Cv14 | 107 | 2.936 | 293.0 | 0.095 | 7.2 | (Das et al., 2016) |
| p27Cv15 | 107 | 2.915 | 293.0 | 0.095 | 7.2 | (Das et al., 2016) |
| p27Cv31 | 107 | 2.81 | 293.0 | 0.095 | 7.2 | (Das et al., 2016) |
| p27Cv44 | 107 | 2.492 | 293.0 | 0.095 | 7.2 | (Das et al., 2016) |
| PTMA | 111 | 3.7 | 288.0 | 0.16 | 7.4 | (Pesce et al., 2023) |
| p15PAF | 111 | 2.81 | 298.0 | 0.15 | 7.0 | (De Biasio et al., 2014) |
| GON7 | 114 | 3.18 | 283.0 | 0.211 | 6.5 | (Arrondel et al., 2019) |
| HAdV5E1A | 114 | 3.6 | 283.0 | 0.24 | 7.0 | (Gonzalez-Foutel et al., 2022) |
| NHE6cmd | 116 | 3.2 | 288.0 | 0.17 | 7.4 | (Pesce et al., 2023) |
| hNL3cyt | 119 | 3.15 | 293.0 | 0.3 | 8.5 | (Paz et al., 2008) |
| hKISS1 | 120 | 3.47 | 283.15 | 0.159 | 7.0 | (Ibáñez de Opakua et al., 2017) |
| RNaseA | 124 | 3.36 | 298.0 | 0.15 | 7.5 | (Riback et al., 2017) |
| hNHE1cdt | 130 | 3.7 | 278.0 | 0.2 | 7.4 | (Kjaergaard et al., 2010) |
| A1S@S150 | 131 | 2.65 | 293.0 | 0.15 | 7.5 | (Martin et al., 2021) |
| HeV_PNT3_3Y3A | 133 | 3.95 | 293.0 | 0.12 | 7.2 | (Nilsson et al., 2022) |
| HeV_PNT3_WT | 133 | 3.67 | 293.0 | 0.12 | 7.2 | (Nilsson et al., 2022) |
| S4L | 134 | 3.6 | 283.15 | 0.169 | 7.2 | (Gomes et al., 2021) |
| NiV_PNT3_WT | 137 | 3.74 | 293.0 | 0.12 | 7.2 | (Nilsson et al., 2022) |
| -10R+10K | 137 | 2.85 | 298.0 | 0.15 | 7.0 | (Bremer et al., 2022) |
| -6R+6K | 137 | 2.79 | 298.0 | 0.15 | 7.0 | (Bremer et al., 2022) |
| +7R | 137 | 2.71 | 298.0 | 0.15 | 7.0 | (Bremer et al., 2022) |
| +2R | 137 | 2.62 | 298.0 | 0.15 | 7.0 | (Bremer et al., 2022) |
| -6R | 137 | 2.57 | 298.0 | 0.15 | 7.0 | (Bremer et al., 2022) |
| -10R | 137 | 2.67 | 298.0 | 0.15 | 7.0 | (Bremer et al., 2022) |
| -9F+3Y | 137 | 2.68 | 298.0 | 0.15 | 7.0 | (Bremer et al., 2022) |
| -8F+4Y | 137 | 2.71 | 298.0 | 0.15 | 7.0 | (Bremer et al., 2022) |
| -9F+6Y | 137 | 2.66 | 298.0 | 0.15 | 7.0 | (Bremer et al., 2022) |
| +7F-7Y | 137 | 2.72 | 298.0 | 0.15 | 7.0 | (Bremer et al., 2022) |
| -12F+12Y | 137 | 2.6 | 298.0 | 0.15 | 7.0 | (Bremer et al., 2022) |
| A1 | 137 | 2.76 | 298.0 | 0.15 | 7.0 | (Bremer et al., 2022) |
| -4D | 137 | 2.64 | 298.0 | 0.15 | 7.0 | (Bremer et al., 2022) |
| +4D | 137 | 2.72 | 298.0 | 0.15 | 7.0 | (Bremer et al., 2022) |
| -3R+3K | 137 | 2.63 | 298.0 | 0.15 | 7.0 | (Bremer et al., 2022) |
| +12D | 137 | 2.8 | 298.0 | 0.15 | 7.0 | (Bremer et al., 2022) |
| +12E | 137 | 2.85 | 298.0 | 0.15 | 7.0 | (Bremer et al., 2022) |
| +7K+12D | 137 | 2.92 | 298.0 | 0.15 | 7.0 | (Bremer et al., 2022) |
| +7K+12Db | 137 | 2.56 | 298.0 | 0.15 | 7.0 | (Bremer et al., 2022) |
| -12F+12Y-10R | 137 | 2.61 | 298.0 | 0.15 | 7.0 | (Bremer et al., 2022) |
| -10F+7R+12D | 137 | 2.86 | 298.0 | 0.15 | 7.0 | (Bremer et al., 2022) |
| +8D | 137 | 2.69 | 298.0 | 0.15 | 7.0 | (Bremer et al., 2022) |
| aSyn140 | 140 | 3.55 | 293.0 | 0.2 | 7.4 | (Ahmed et al., 2021) |
| TtASR1 | 141 | 3.31 | 293.15 | 0.15 | 7.3 | (Hamdi et al., 2017) |
| P300 | 141 | 3.3 | 293.0 | 0.17 | 7.5 | (Borysik et al., 2015) |
| Calpastatin | 141 | 3.9 | 293.0 | 0.17 | 7.5 | (Borysik et al., 2015) |
| HvASR1 | 143 | 3.51 | 293.15 | 0.15 | 7.3 | (Hamdi et al., 2017) |
| FhuA | 144 | 3.34 | 298.0 | 0.15 | 7.5 | (Riback et al., 2017) |
| TIF2NRID | 150 | 3.74 | 283.15 | 0.175 | 6.8 | (Senicourt et al., 2021) |
| ED2 | 154 | 4.13 | 293.15 | 0.153 | 7.4 | (Gondelaud et al., 2021) |
| ED4 | 163 | 4.06 | 293.15 | 0.153 | 7.4 | (Gondelaud et al., 2021) |
| AavLEA1 | 163 | 4.1 | 293.0 | 0.17 | 7.5 | (Borysik et al., 2015) |
| ANAC046 | 167 | 3.6 | 298.0 | 0.14 | 7.0 | (Pesce et al., 2023) |
| K27 | 167 | 3.7 | 288.0 | 0.15 | 7.4 | (Mylonas et al., 2008) |
| K10 | 168 | 4.0 | 288.0 | 0.15 | 7.4 | (Mylonas et al., 2008) |
| PARCL | 180 | 3.43 | 293.15 | 0.17 | 7.5 | (Ostendorp et al., 2022) |
| ERD14 | 184 | 4.6 | 293.0 | 0.17 | 7.5 | (Borysik et al., 2015) |
| K25 | 185 | 4.1 | 288.0 | 0.15 | 7.4 | (Mylonas et al., 2008) |
| N_FATZ1 | 191 | 3.45 | 293.15 | 0.192 | 7.5 | (Sponga et al., 2021) |
| K32 | 198 | 4.2 | 288.0 | 0.15 | 7.4 | (Mylonas et al., 2008) |
| D91_FATZ1 | 209 | 3.86 | 293.15 | 0.192 | 7.5 | (Sponga et al., 2021) |
| MetC | 217 | 5.16 | 288.15 | 0.167 | 7.5 | (Kolonko et al., 2016) |
| cDAXX | 246 | 4.75 | 293.0 | 0.13 | 8.0 | (Schmit et al., 2019) |
| K23 | 254 | 4.9 | 288.0 | 0.15 | 7.4 | (Mylonas et al., 2008) |
| RNaseE603_850 | 254 | 5.26 | 288.0 | 0.274 | 7.5 | (Bruce et al., 2018) |
| tau35 | 255 | 4.64 | 293.2 | 0.15 | 7.4 | (Lyu et al., 2021) |
| ERD10 | 259 | 6.0 | 293.0 | 0.17 | 7.5 | (Borysik et al., 2015) |
| ED1 | 260 | 5.42 | 293.15 | 0.153 | 7.4 | (Gondelaud et al., 2021) |
| CoRNID | 271 | 4.7 | 293.15 | 0.192 | 7.5 | (Cordeiro et al., 2019) |
| K44 | 283 | 5.2 | 288.0 | 0.15 | 7.4 | (Mylonas et al., 2008) |
| PNtdeltaPG | 310 | 4.99 | 293.15 | 0.175 | 7.5 | (Baxa et al., 2024) |
| Msh6_NTR | 313 | 5.6 | 293.0 | 0.12 | 8.0 | (Shell et al., 2007) |
| PNtS5 | 334 | 4.87 | 298.0 | 0.15 | 7.5 | (Bowman et al., 2020) |
| PNtS4 | 334 | 5.34 | 298.0 | 0.15 | 7.5 | (Bowman et al., 2020) |
| PNtS1 | 334 | 4.92 | 298.0 | 0.15 | 7.5 | (Bowman et al., 2020) |
| PNt | 334 | 5.11 | 298.0 | 0.15 | 7.5 | (Bowman et al., 2020; Riback et al., 2020) |
| PNtS6 | 334 | 5.26 | 298.0 | 0.15 | 7.5 | (Bowman et al., 2020) |
| GHRICD | 351 | 6.0 | 298.0 | 0.35 | 7.3 | (Seiffert et al., 2020; Pesce et al., 2023) |
| ED3 | 373 | 6.51 | 293.15 | 0.153 | 7.4 | (Gondelaud et al., 2021) |
| ht2N3R | 410 | 6.3 | 293.2 | 0.15 | 7.4 | (Lyu et al., 2021) |
| ht2N4R | 441 | 6.7 | 293.2 | 0.15 | 7.4 | (Lyu et al., 2021) |
| MAP2c | 467 | 6.6 | 293.0 | 0.16 | 6.9 | (Melková et al., 2018) |

**Table S2.** Experimental solution conditions and radii of gyration of the disordered proteins included in the validation set. Errors of experimental *R*_g_ are estimated as 1% of *R*_g_ values.

| Protein | N | $R_g$ [nm] | T [K] | $c_s$ [M] | pH | Ref. |
| --- | --- | --- | --- | --- | --- | --- |
| Nup49_Rg | 38 | 1.6 | 296.0 | 0.15 | 7.4 | (Fuertes et al., 2017) |
| NLS_Rg | 46 | 2.4 | 296.0 | 0.15 | 7.4 | (Fuertes et al., 2017) |
| dChMINUS_Rg | 68 | 2.58 | 295.0 | 0.172 | 7.3 | (Holla et al., 2024) |
| sNhPLUS_Rg | 68 | 2.55 | 295.0 | 0.172 | 7.3 | (Holla et al., 2024) |
| Red1 | 70 | 2.43 | 293.15 | 0.157 | 7.5 | (Soni et al., 2023) |
| HMPVP_NTD | 80 | 2.74 | 293.0 | 0.17 | 7.5 | (Renner et al., 2017) |
| NUS_Rg | 82 | 2.5 | 296.0 | 0.15 | 7.4 | (Fuertes et al., 2017) |
| SMAD2_linker | 90 | 2.99 | 283.0 | 0.15 | 7.2 | (Gomes et al., 2021) |
| IBB_Rg | 99 | 3.2 | 296.0 | 0.15 | 7.4 | (Fuertes et al., 2017) |
| NUL_Rg | 114 | 3.0 | 296.0 | 0.15 | 7.4 | (Fuertes et al., 2017) |
| TRPV4 | 133 | 3.2 | 293.15 | 0.118 | 7.0 | (Goretzki et al., 2023) |
| V2 | 137 | 2.31 | 298.0 | 0.15 | 7.0 | (Pesce et al., 2024) |
| V5 | 137 | 2.48 | 298.0 | 0.15 | 7.0 | (Pesce et al., 2024) |
| V4 | 137 | 2.39 | 298.0 | 0.15 | 7.0 | (Pesce et al., 2024) |
| V3 | 137 | 2.35 | 298.0 | 0.15 | 7.0 | (Pesce et al., 2024) |
| CTR_XRCC4 | 138 | 3.42 | 298.0 | 0.16 | 6.5 | (Vu et al., 2024) |
| N98_Rg | 153 | 2.9 | 296.0 | 0.15 | 7.4 | (Fuertes et al., 2017) |
| CTir | 173 | 3.81 | 298.15 | 0.179 | 6.5 | (Vieira et al., 2024) |
| Nsp1_Rg | 178 | 4.1 | 296.0 | 0.15 | 7.4 | (Fuertes et al., 2017) |
| ER_NTD | 187 | 3.07 | 277.15 | 0.198 | 7.4 | (Du et al., 2025) |

**Table S3.** Experimental solution conditions and PRE data included in the validation set.

| Protein | N | $N_{labels}$ | $\omega_I/2\pi [MHz]$ | T [K] | $c_s$ [M] | pH | Ref. |
| --- | --- | --- | --- | --- | --- | --- | --- |
| aSyn | 140 | 5 | 700 | 283 | 0.2 | 7.4 | (Dedmon et al., 2005) |
| OPN | 220 | 10 | 800 | 298 | 0.15 | 6.5 | (Kurzbach et al., 2016) |
| FUS | 163 | 3 | 850 | 298 | 0.15 | 5.5 | (Monahan et al., 2017) |
| FUS12E | 164 | 3 | 850 | 298 | 0.15 | 5.5 | (Monahan et al., 2017) |

**Table S4.** Experimental solution conditions and FRET data included in the validation set.

| Protein | N | FRET efficiency ± Error | T [K] | $c_s$ [M] | pH | Ref. |
| --- | --- | --- | --- | --- | --- | --- |
| sGrich | 68 | $0.861 \pm 0.008$ | 295.0 | 0.172 | 7.3 | (Holla et al., 2024) |
| dkh | 68 | $0.77 \pm 0.02$ | 295.0 | 0.172 | 7.3 | (Holla et al., 2024) |
| dGrich | 68 | $0.71 \pm 0.02$ | 295.0 | 0.172 | 7.3 | (Holla et al., 2024) |
| sPTBP | 68 | $0.66 \pm 0.02$ | 295.0 | 0.172 | 7.3 | (Holla et al., 2024) |
| sCh | 68 | $0.63 \pm 0.04$ | 295.0 | 0.172 | 7.3 | (Holla et al., 2024) |
| sNrich | 68 | $0.61 \pm 0.02$ | 295.0 | 0.172 | 7.3 | (Holla et al., 2024) |
| skh | 68 | $0.595 \pm 0.005$ | 295.0 | 0.172 | 7.3 | (Holla et al., 2024) |
| dChPLUS | 68 | $0.59 \pm 0.03$ | 295.0 | 0.172 | 7.3 | (Holla et al., 2024) |
| dChMINUS | 68 | $0.5 \pm 0.03$ | 295.0 | 0.172 | 7.3 | (Holla et al., 2024) |
| skl | 68 | $0.49 \pm 0.03$ | 295.0 | 0.172 | 7.3 | (Holla et al., 2024) |
| sNhPLUS | 68 | $0.49 \pm 0.03$ | 295.0 | 0.172 | 7.3 | (Holla et al., 2024) |
| dArich | 68 | $0.48 \pm 0.02$ | 295.0 | 0.172 | 7.3 | (Holla et al., 2024) |
| dErich | 68 | $0.47 \pm 0.03$ | 295.0 | 0.172 | 7.3 | (Holla et al., 2024) |
| dTRBP | 68 | $0.45 \pm 0.03$ | 295.0 | 0.172 | 7.3 | (Holla et al., 2024) |
| dki | 68 | $0.43 \pm 0.03$ | 295.0 | 0.172 | 7.3 | (Holla et al., 2024) |
| sNhMINUS | 68 | $0.42 \pm 0.03$ | 295.0 | 0.172 | 7.3 | (Holla et al., 2024) |
| Nup49 | 38 | $0.87 \pm 0.02$ | 296.0 | 0.15 | 7.4 | (Fuertes et al., 2017) |
| NLS | 46 | $0.79 \pm 0.02$ | 296.0 | 0.15 | 7.4 | (Fuertes et al., 2017) |
| NUS | 82 | $0.53 \pm 0.02$ | 296.0 | 0.15 | 7.4 | (Fuertes et al., 2017) |
| IBB | 99 | $0.5 \pm 0.02$ | 296.0 | 0.15 | 7.4 | (Fuertes et al., 2017) |
| NUL | 114 | $0.48 \pm 0.02$ | 296.0 | 0.15 | 7.4 | (Fuertes et al., 2017) |
| Sic1 | 92 | $0.42 \pm 0.02$ | 293.0 | 0.2 | 7.5 | (Gomes et al., 2020) |

**Table S5.**
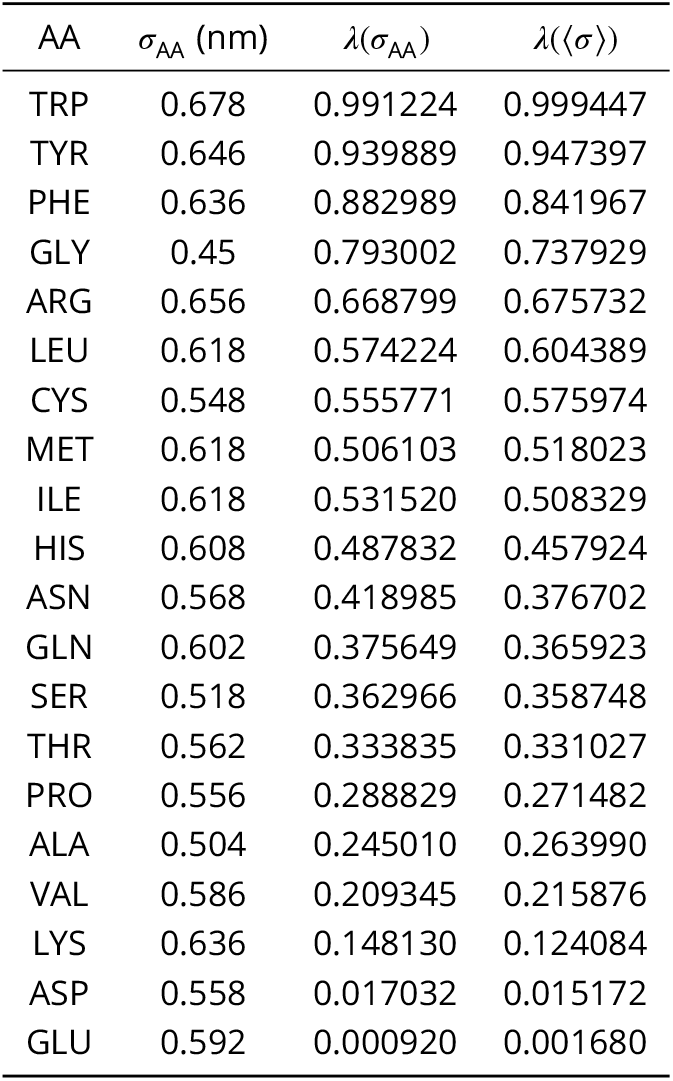
Amino acid specific stickiness parameters. *σ*_AA_ represents the amino acid specific radii (***Kim and Hummer, 2008***), and *λ*(*σ*_AA_) are the stickiness values obtained using these radii. *λ*(⟨*σ*⟩) are the values obtained using a shared, composition-weighted average value ⟨*σ*⟩ = 0.56 nm.

| AA | $\sigma_{AA}$ (nm) | $\lambda(\sigma_{AA})$ | $\lambda(\langle\sigma\rangle)$ |
| --- | --- | --- | --- |
| TRP | 0.678 | 0.991224 | 0.999447 |
| TYR | 0.646 | 0.939889 | 0.947397 |
| PHE | 0.636 | 0.882989 | 0.841967 |
| GLY | 0.45 | 0.793002 | 0.737929 |
| ARG | 0.656 | 0.668799 | 0.675732 |
| LEU | 0.618 | 0.574224 | 0.604389 |
| CYS | 0.548 | 0.555771 | 0.575974 |
| MET | 0.618 | 0.506103 | 0.518023 |
| ILE | 0.618 | 0.531520 | 0.508329 |
| HIS | 0.608 | 0.487832 | 0.457924 |
| ASN | 0.568 | 0.418985 | 0.376702 |
| GLN | 0.602 | 0.375649 | 0.365923 |
| SER | 0.518 | 0.362966 | 0.358748 |
| THR | 0.562 | 0.333835 | 0.331027 |
| PRO | 0.556 | 0.288829 | 0.271482 |
| ALA | 0.504 | 0.245010 | 0.263990 |
| VAL | 0.586 | 0.209345 | 0.215876 |
| LYS | 0.636 | 0.148130 | 0.124084 |
| ASP | 0.558 | 0.017032 | 0.015172 |
| GLU | 0.592 | 0.000920 | 0.001680 |

